# The optimization of *Salmonella* surveillance programmes for pullet and layer farms using local farm density as a risk factor

**DOI:** 10.1101/2023.09.10.557094

**Authors:** Peter H.F. Hobbelen, Thomas J. Hagenaars, Michal Peri Markovich, Michel Bellaiche, Armando Giovannini, Fabrizio De Massis, Aline de Koeijer

## Abstract

Human salmonellosis cases are often caused by Salmonella serovars Enteritidis and Typhimurium and a large percentage of Salmonella outbreaks is associated with the consumption of eggs and egg products. For this reason, many countries implemented general surveillance programmes for the detection and control of *Salmonella* on pullet and layer farms. The infection risk however varies between farms and the identification of risk factors for *Salmonella* infection may be used to improve the performance of these surveillance programmes. The aims of this study are therefore to determine 1) whether local farm density is a risk factor for the infection of pullet and layer farms by *Salmonella* Enteritidis and Typhimurium and 2) whether the sampling effort of surveillance programmes can be reduced by accounting for this risk factor, while still providing sufficient control of target serovars Enteritidis and Typhimurium. To assess the importance of local farm density as a risk factor, we fitted different transmission kernels to Israeli surveillance data during the period from June 2017 to April 2019. The analysis showed that the distance to infected farms significantly increases the infection risk by serovar Enteritidis within an approximately 4 km radius and by Typhimurium within an approximately 0.3 km radius. We subsequently used these kernels to derive a model for the between-farm R_0_ and used it to optimise a surveillance programme that subdivided layer farms into groups at low and at high risk of between-farm transmission based on the local farm density and allowed the sampling frequency to vary between these groups. In this design, the pullet farms were always sampled one week prior to pullet distribution. Our analysis showed that the risk-based surveillance programme was able to keep the between-farm R_0_ of serovars Enteritidis and Typhimurium below 1 for all pullet and layer farms, using a sampling effort that was reduced by 32% compared to the currently implemented surveillance programme in Israel. The results of our study therefore indicate that local farm density is an important risk factor for infection of pullet and layer farms by *Salmonella* Enteritridis and Typhimurium and can be used to improve the performance of surveillance programmes.

## Introduction

Salmonellosis is globally one of the most common foodborne diseases [1, 2]. Typical symptoms are diarrhoea, stomach cramps and fever[3]. Although most cases are mild, severe and sometimes life-threatening complications can develop[4]. The genus *Salmonella* contains only two species, but these can be further subdivided into more than 2600 serovars with varying epidemiological characteristics such as the severity of symptoms, the risk of complications and resistance to antibiotics[5]. Human *Salmonella* infections are primarily caused by the consumption of contaminated food products such as raw or undercooked meat, eggs, dairy products and vegetables[6], although other transmission pathways such as contact with animals and human to human transmission are possible, too.

The consumption of eggs and egg products is responsible for a large part of the human *Salmonella* cases. According to a recent estimate for example, around 45% of reported foodborne *Salmonella* outbreaks in the European Union with strong evidence for the source of infection were attributed to the consumption of eggs and egg products[2]. For this reason, many developed countries introduced legislation to monitor and control the prevalence of *Salmonella* during the pre-harvest, processing and post-harvest stages of the egg (product) supply chain. Since the egg supply chain has a pyramid shape with more animals and eggs involved at every subsequent stage[7], control measures that reduce the prevalence of *Salmonella* early in the supply chain will potentially be most efficient. One of these measures is the implementation of a *Salmonella* surveillance programme for pullet and layer farms[8]. The investment that is needed to implement a surveillance programme depends amongst others on the sampling effort that is required to achieve the goals of such a programme. Currently, mandatory *Salmonella* surveillance programmes of pullet and layer farms in developed countries often use the same sampling interval for all layer farms[8]. The performance of these programmes may be improved by adjusting the sampling effort of farms based on their infection risk by *Salmonella*[9]. The design of such a risk-based surveillance scheme requires knowledge on factors that 1) have a significant influence on the risk of *Salmonella* infection for pullet and layer farms, 2) are relatively persistent in time and 3) vary between farms within the surveillance area.

It may be hypothesized that pullet and layer farms can become infected by *Salmonella* as a result of between-farm transmission in addition to introduction from a source in the local environment. If this is the case, potential risk factors of *Salmonella* infection of pullet and layer farms may be derived from the relationship between the infection probability and the distance to infected farms. Literature on this subject is however scarce. There are a few studies that included the distance to the nearest farm in a risk factor analysis for the infection probability of layer farms with *Salmonella*. These studies treated between-farm distance as a categorical variable using a threshold distance of 0.5 or 1 km[10–12]. This implicitly assumes a sharp drop in the infection probability above a certain distance between farms. Another paper looked at the relationship between the density of farms and the occurrence of layer farms infected with *Salmonella* without reporting the significance of this effect[13]. This paper used a 1 km^2^ area to calculate the farm density, but did not provide arguments to support this choice. However, the relevance of farm density as a risk factor may only appear when the spatial range used to calculate the density is representative for the range within which between-farm transmission is possible. Information on the precise form of spatial kernels describing between-farm transmission would therefore be useful to underpin the definition of more relevant and significant risk factors. Such spatial transmission kernels can also be used to evaluate disease control strategies, for example by calculating the between-farm reproduction number[14] (R_0_) or by incorporating them into spatial simulation models that describe the infection dynamics of *Salmonella* at the farm level[15]. More detailed information about the form of transmission kernels may therefore also be useful for the design of surveillance programmes that aim to provide sufficient control of *Salmonella* target serovars. To our knowledge, no study tried to formulate transmission kernels describing the infection probability of *Salmonella* as a function of the distance between infected and susceptible farms before.

Israel implemented a general surveillance programme for *Salmonella* on pullet and layer farms in 2017. The aim of the programme was to identify and monitor the prevalence of *Salmonella* serovars and to reduce the prevalence of the target serovars Enteritidis and Typhimurium, that are often associated with human Salmonellosis cases[2]. Layer and pullet farms in Israel may be clustered together in small villages, but can also lie at relatively isolated locations. As a result, there is a much variation in the distance to neighbouring farms and the local farm density. If between-farm transmission plays an important role in the disease dynamics for this system, it may be possible to reduce the sampling effort by sampling farms in areas with a high farm density more frequently than farms in low density areas.

Given the host-pathogen system consisting of pullet and layer farms in Israel and *Salmonella* target serovars Enteritidis and Typhimurium, the aims of this study are therefore to 1) determine the significance of between-farm transmission for the infection dynamics at the farm level, 2) explore the relationship between the infection probability of a farm and its distance to infected farms and 3) use this relationship to design and evaluate a risk-based surveillance programme that provides sufficient disease control of *Salmonella* target serovars.

## Materials and Methods

### The current *Salmonella* surveillance programme for layer and pullet farms in Israel

Under the present surveillance scheme for *Salmonella* that started in June 2017, layer farms are sampled every 15 weeks from the moment the flock reaches an age of 22 to 24 weeks. Pullet farms are sampled approximately one week prior to the distribution of pullets to layer farms. Samples were taken by inspectors from either the Israeli Veterinary Services (IVS) or the Egg and Poultry Board (EPB). On each sampling occasion, IVS inspectors usually took 2 pairs of drag swabs, 2 pairs of dust swabs from walls and equipment and one dust sample, while EPB inspectors usually took 2 pairs of drag swabs. If present, *Salmonella* serovars were identified to the group level according to the Kauffman-White scheme by the EPB laboratory. Isolates of groups B or D were further identified to the serovar level by the *Salmonella* National Reference Laboratory of the Israeli Ministry of Health. Farms testing positive for *Salmonella* serovars Enteritidis and/or Typhimurium were culled and thoroughly cleaned and disinfected. Repopulation of these farms was only allowed after drag swabs from the empty houses tested negative for all *Salmonella* serovars. Hereafter, we will refer to *Salmonella* serovars Enteritidis and Typhimurium simply as SE and ST, respectively.

### Data

Sampling data from the *Salmonella* surveillance scheme were available for a total of 1,842 layer farms and 56 pullet farms during the period from early June 2017 to early April 2019. Together, the layer and pullet farms made up around 60% of all poultry farms in Israel during this period. Hens were caged on the vast majority of layer farms (97.5%) and free-ranging on the remainder of farms. Farm size varied widely (less than 100 to 255,000), and the median size was around 2,900. The number of houses per farm varied from 1 to 6, but the vast majority of farms contained only one house (97.9%). For farms that had several houses, the production cycle always started at the same date in all houses. Production cycles on farms that did not test positive for SE or ST lasted on average 18.2 weeks for pullets and 95.4 weeks for layers.

### Reconstruction of the disease status of farms

The original data were organised at the level of individual samples with each row containing information on amongst others the sampling date and type, the test result and specifics of the sampled farm. For the analysis of the importance of local farm density as a risk factor for disease transmission below, the information on individual samples must be combined in order to reconstruct the disease status of individual farms in time. The disease status of a farm was reconstructed separately for SE as well as ST and indicated whether a farm was 1) susceptible to infection, 2) infectious to other farms or 3) empty. It was assumed that empty farms were neither susceptible nor infectious. In order to reconstruct the disease status of an individual farm, it is necessary to have information on the starting and end date of subsequent production cycles, the sampling dates and test results. In addition, the location of each farm should be available to perform the analysis below. This type of information was sometimes missing or data on for example the start and end dates of consecutive production cycles on the same farm were conflicting. We therefore developed a protocol that describes how we dealt with missing data and conflicting information (S1 Appendix, sections 1 and 2). In the end, we were able to reconstruct the disease status during the period from June 2017 to April 2019 for 96.6% of farms in the dataset and the remainder of farms were excluded from the analysis due to incomplete or conflicting data. We estimated the day that a farm became infected with SE or ST as the date in the middle of the interval bounded by the last negative and first positive sample. If the first sample of a production cycle was immediately positive, the date of infection was determined as described in S1 Appendix (section 1). We assumed that farms were infectious from the estimated day of infection until the end of the production cycle (usually due to culling). After the analyses described in this paper were completed, a few additional flocks were found to be positive for SE and ST during the study period for a variety of reasons (S1 Appendix, section 3). This number of flocks was small compared to the total number of positive flocks that was included in our analysis and we therefore assumed that the outcomes of the analyses would not have substantially changed by including the new information.

### Local farm density as risk factor for *Salmonella* infection

To determine whether the local farm density is a risk factor for the transmission of SE and/or ST between farms, we formulated the infection probability of a susceptible farm as a function of the distance to an infectious farm. Assuming that the number of infection events per unit of time follows a Poisson distribution, the probability (*P*_inf,i_) that a susceptible farm *i* becomes infected during time interval *T* can be calculated as

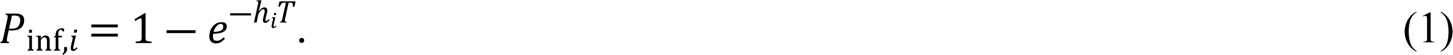

We formulated the hazard function *h*_i_ as a sum of two terms

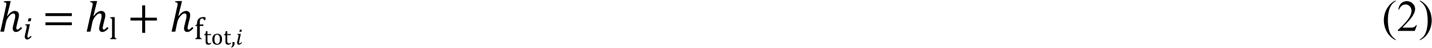

with *h*_1_ representing transmission from a local source in the environment to the susceptible farm and ℎ_ftot,*i*_ representing between-farm transmission. For clarity, we defined a local source in the environment of a given farm *i* as a source that also exists in the absence of other farms in the surroundings of farm *i*. Hereafter, we will simplify the term “local source in the environment” to “local source”. We assumed that the hazard of infection from a local source per unit of time was the same for all farms. The hazard of infection of susceptible farm *i* by all infectious farms together per unit of time was calculated as

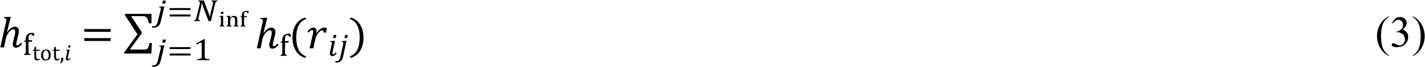

with *N*_inf_ denoting the total number of farms that was infectious at a certain point in time and *r*_ij_ denoting the distance between susceptible farm *i* and infectious farm *j* (S1 Appendix, section 4). Function *h*_f_ describes the hazard of infection of a susceptible farm by an infectious farm per unit of time as a function of the distance between these farms. We compared two hazard functions (*h*_i_) to assess the importance of between-farm distance as a risk factor for infection by SE and ST. The first function assumes that local sources are the only source of infection, i.e. ℎ_ftot,*i*_= 0. The second function assumes that transmission from local sources as well as between-farm transmission are possible, with the hazard of infection of farm *i* by farm *j* per unit of time formulated as a step function

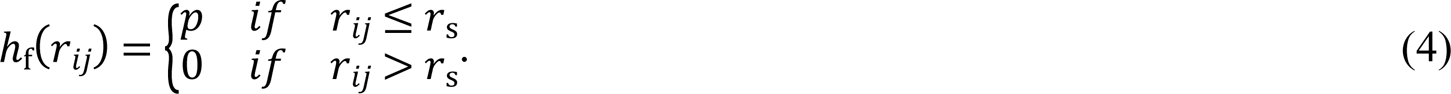

This function assumes that the hazard of between-farm transmission is constant up to a certain threshold distance *r*_s_ and then sharply decreases to zero for larger distances. We parameterized these two hazard functions by fitting them to the data on the disease status of farms in time using a maximum likelihood approach[14]. We determined 95% confidence intervals for kernel parameters using the likelihood ratio test[16]. The most parsimonious hazard function was determined using Akaike’s Information Criterion (AIC). If the constant hazard function has a significantly higher AIC value than the step function (less parsimonious), this suggests that between-farm transmission contributes to the infection probability when the distance to infected farms remains below a certain threshold. In turn, this implies that the local farm density may be a risk factor for infection by Salmonella. We performed separate analyses for SE and ST.

### The dependence of the between-farm R_0_ on the sampling interval of a surveillance programme

In this study, we used the local between-farm R_0_ to evaluate and optimize different surveillance programmes (see below). The between-farm R_0_ for a given infectious farm *j* is defined as the number of farms that on average will become infected by this farm, assuming that all other farms are susceptible. This R_0_ can be calculated as the sum of the infection probabilities of the individual susceptible farms as

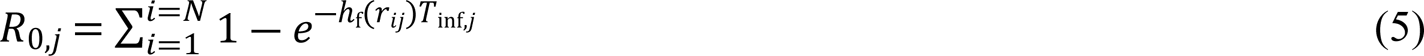

with *N* representing the total number of farms and *T*_inf,j_ denoting the infectious period of farm *ij*. In this study, the total number of farms *N* consisted of all layer and pullet farms in Israel for which the disease status in time could be reconstructed.

As can be seen from Equation 5, the R_0_ of a given farm depends on its infectious period and therefore on the length of the sampling interval for this farm, because farms will be culled after testing positive for SE and/or ST. This link between the between-farm R_0_ and the length of the sampling interval can be used to evaluate and optimize surveillance programmes.

Assuming that farms become infected on average in the middle of the period between two sampling events, the infectious period (*T*_inf_) can be derived from the length of the sampling interval (*T*_sample_) as

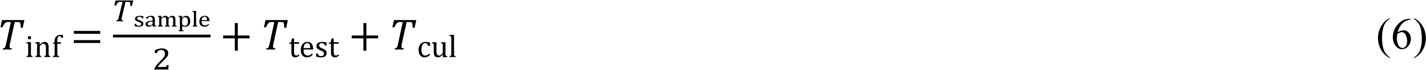

with *T*_test_ denoting the time interval between the sampling date and the date on which the test result became available and *T*_cul_ denoting the time interval between the latter date and the culling of the farm. Both these intervals were estimated to be 1 week long.

Hereafter, we will refer to the between-farm R_0_ simply as R_0_.

### Surveillance programmes

In this study, we analysed the performance of different types of surveillance programmes for three serovar scenarios with scenario A focusing on the monitoring and control of SE, scenario B focusing on ST and scenario C focusing on both these *Salmonella* serovars. For all serovar scenarios, we analysed the performance of two types of surveillance programmes. Firstly, a general surveillance programme with one common sampling interval for all layer farms. Secondly, a surveillance programme that accounts for the local farm density as a risk factor. This was done by assigning each layer farm into either a low-risk or a high-risk group, based on the local farm density. The local farm density for a given layer farm *i* was calculated as the total number of farms in a circular area with farm *i* at the centre. The radius of the circle was set to the distance *r*_s_ in the step function describing the hazard of between-farm transmission, as this distance models the range within which this hazard is relevant (Equation 4). If the local farm density was below a certain threshold, farm *i* was assigned to the low-risk group and otherwise to the high-risk group. The sampling interval of layer farms was allowed to vary between the low-risk and high-risk group. For both the general and risk-based surveillance programme, we assumed that layer farms should at least be sampled once a year. In addition, for both surveillance programmes we assumed that pullet farms were sampled one week before the end of the production cycle (sampling time at flock age 17 wks) in order to prevent the distribution of infected pullets to layer farms.

### The criterion for comparing surveillance programmes

We compared the performance of different surveillance programmes for a given serovar scenario based on the minimum number of sampling events that kept the R_0_ of all serovars included in the surveillance programme < 1, for all layer and pullet farms during a time period of one year. Here a visit to a farm by an inspector is counted as one sampling event, independent of the number and types of samples taken during that visit.

### The optimization of surveillance programmes

For the general surveillance programme, we used the following algorithm to determine the optimum sampling interval for layer farms. Firstly, in order to find the sampling interval that resulted in R_0_ < 1 for all farms, we identified the farm with the highest R_0_ for a 1 week sampling interval, since this farm will have the highest R_0_ for any sampling interval. Denoting this farm as farm *k*, we subsequently increased the sampling interval using weekly steps and recalculated the R_0_ of farm *k* every step until it became ≥ 1. The sampling interval of all layer farms was then set to the longest sampling interval for which the R_0_ of farm *k* was still lower than 1.

The risk-based surveillance programme is specified by three parameters, namely one threshold density that is used to divide layer farms between the low and high-risk groups and one sampling interval for each risk group. To find the optimum values for these parameters, we first created a sequence of threshold densities ranging from the lowest to the highest local farm density (see above). For serovar scenario A (targeting SE), the local farm density ranged from 0 to approximately 5 farms per km^2^ and the threshold density was varied in steps of 0.05. For serovar scenario B (targeting ST), the local farm density ranged from 0 to approximately 175 farms per km^2^ and the threshold density was varied in steps of 5. The difference in the range of farm densities between both serovars is due to the difference in the radius of the circular area used to calculate the local farm density for each serovar (see above). For each threshold density in the sequence, we determined the corresponding subdivision of layer farms into the low-risk and high-risk groups. For each risk-group, we subsequently applied the algorithm described above to determine the longest sampling interval resulting in R_0_ < 1 for all layer farms in the group. In this way, we obtained the optimum sampling intervals for each threshold density and subsequently derived the total number of sampling events per year for all farms combined from these optimum sampling intervals. The combination of threshold density and sampling intervals that required the lowest number of sampling events per year to keep the R_0_ of all farms <1 was selected as the optimal risk-based surveillance programme.

For serovar scenario C, targeting both serovars, the local farm density was calculated using the threshold distance of the hazard function for SE, since on most farms the R_0_ corresponding to a certain sampling interval was higher for SE than for ST. The algorithm used to optimize the sampling interval for a group of layer farms was the same as described above for serovar scenarios A and B with one exception. In order to find the farm with the highest R_0_, we calculated the R_0_ of all layer farms for SE as well as ST.

## Results

### The number of *Salmonella* Enteritidis and Typhimurium cases in time

Both the number of detected SE and ST cases have dropped steadily during the period for which surveillance data were available (Fig 1). The disease prevalence on layer farms during the first 24 weeks after the start of the current surveillance programme was 2.5% for SE and 1.5% for ST. During the last 24 weeks for which data was available, the prevalence had declined to 0% for SE and 0.1% for ST.

**Fig 1.**
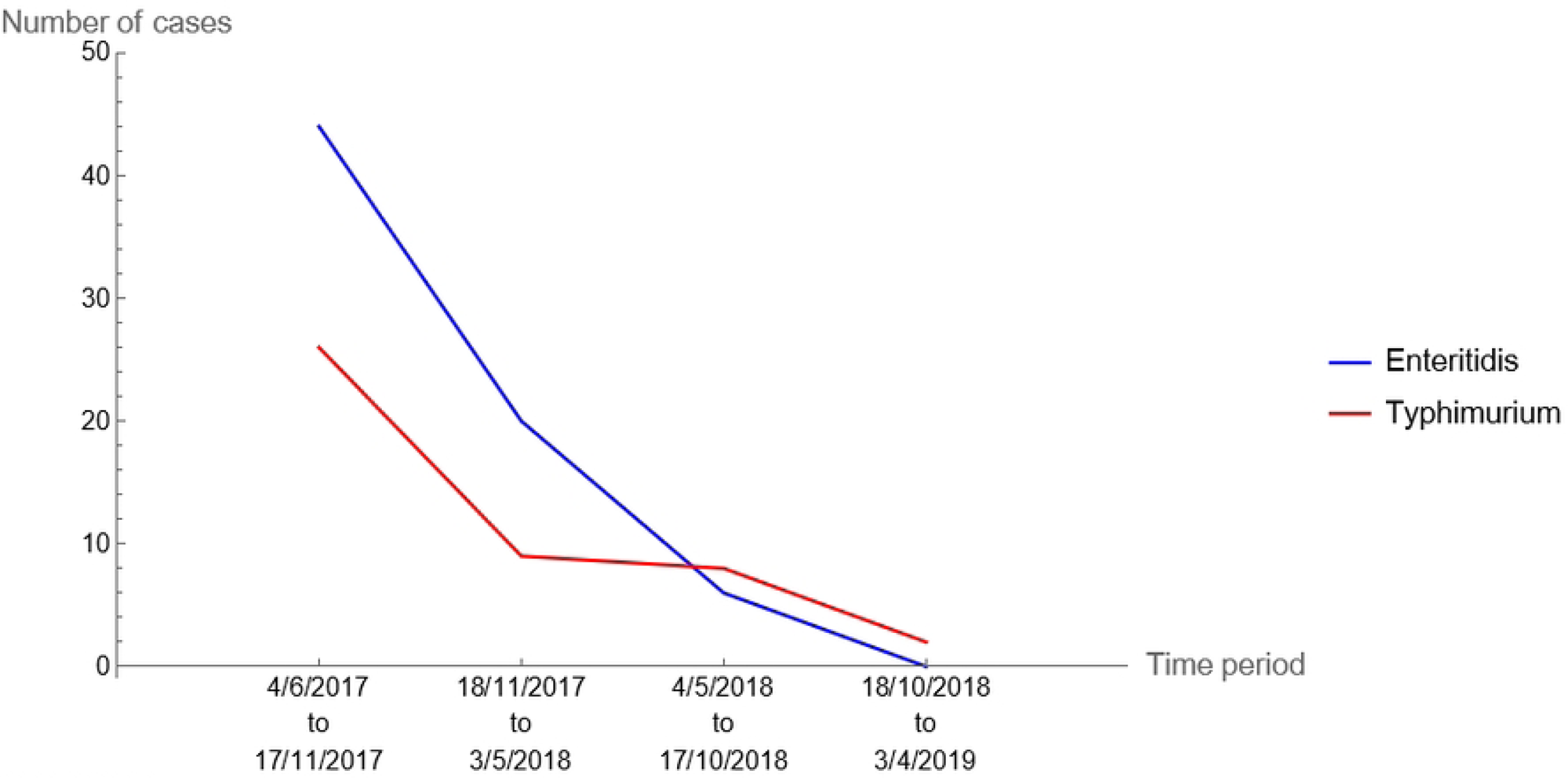
The number of detected *Salmonella* Enteritidis and Typhimurium cases in Israeli pullet and layer farms during the period from early June 2017 to early April 2019, subdivided into 4 consecutive time intervals of approximately 24 weeks (167 days).

### The local farm density as a risk factor for *Salmonella* infection

The hazard functions that account for both introduction from a local source as well as between-farm transmission provided a significantly better fit to the data than the hazard functions that only account for introduction from a local source. This was the case for SE as well as ST (Table 1 and Fig 2) and suggests that the distance to infected farms affects the infection probability of pullet and layer farms in Israel with these *Salmonella* serovars. The local farm density may therefore be a risk factor for Salmonella infection. The hazard of infection was 4.5 times higher when a susceptible farm was located near a farm that was infectious for SE and 6.2 times higher near a farm that was infectious for ST. Interestingly, between-farm transmission of SE seems to occur over much longer distances (up to approximately 4 km) than between-farm transmission of ST (up to approximately 300 m).

**Fig 2.**
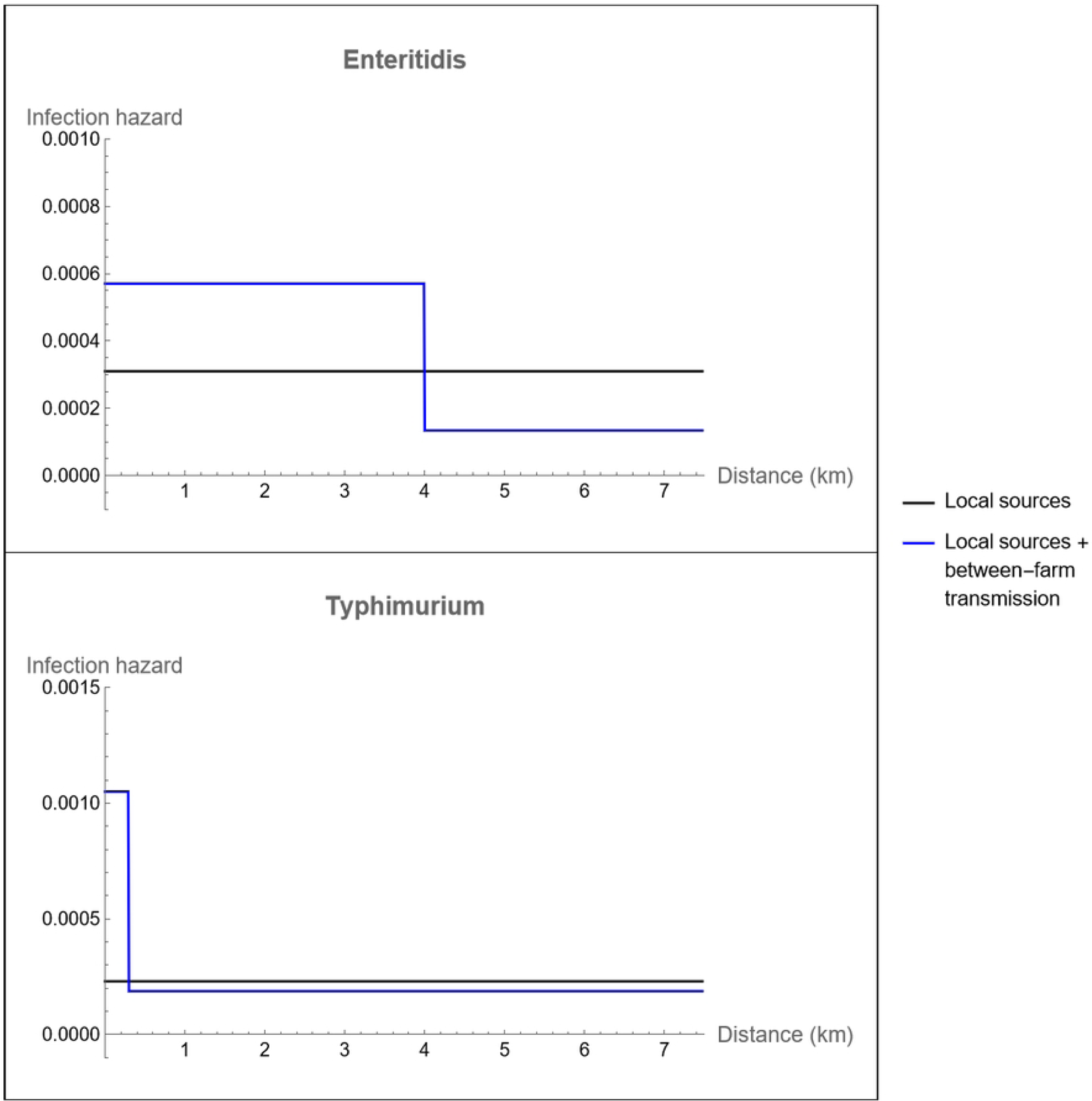
The transmission kernels describing the infection hazard of susceptible layer and pullet farms in Israel due to transmission from a local source in the environment and between-farm transmission for *Salmonella* Enteritidis and Typhimurium.

**Table 1.** Maximum likelihood estimates and 95% confidence intervals (in brackets) for the parameters of the transmission kernels that describe the infection hazard of layer and pullet farms in Israel per unit of time (wks) in terms of both transmission from local sources in the environment as well as and between-farm transmission for *Salmonella* serovars Enteritidis and Typhimurium.

| Hazard function | Hazard due to environmental transmission<sup>a</sup> ($h_e$) | Hazard due to between-farm transmission<sup>b</sup> ($h_f$) | | NLL<sup>c</sup> | AIC<sup>d</sup> |
| --- | --- | --- | --- | --- | --- |
| | | Amplitude ($p$) | Threshold distance ($r_s$) (km) | | |
| *Salmonella* Enteritidis |  |  |  |  |  |
| Only local sources | 3.1e-4 (2.3e-4 - 4.1e-4) | - | - | 435.9 | 873.8 |
| Local sources and between-farm transmission | 1.33e-4 (8e-5 - 2e-4) | 4.63e-4 (3e-4 – 7.2e-4) | 3.91 (3.71 – 4.06) | 405.4 | 816.8 |
| *Salmonella* Typhimurium |  |  |  |  |  |
| Only local sources | 2.3e-4 (1.6e-4 – 3e-4) | - | - | 328.9 | 659.8 |
| Local sources and between-farm transmission | 1.87e-4 (1.3e-4 – 2.6e-4) | 9.76e-4 (3e-4 – 2.3e-3) | 0.27 (0.19 – 0.59) | 322.1 | 650.6 |

<sup>a)</sup> See equation 2 in the main text
<sup>b)</sup> See equation 4 in the main text
<sup>c)</sup> NLL denotes the negative log likelihood
<sup>d)</sup> AIC denotes Akaike's Information Criterion

### R_0_ in the absence of surveillance

The R_0_ value for SE in the absence of a surveillance programme was predicted to be equal or higher than 1 for approximately 74% of the layer farms with a maximum value of 5.4 (S1 Fig). For ST, this percentage was smaller (approximately 19%) and the highest R_0_ value amounted to 2.2 (S2 Fig). This suggests that between-farm transmission is an important pathway for *Salmonella* infection in the absence of disease control. Interestingly, the R_0_ value of SE as well as of ST were far below 1 for pullet farms and amounted to 0.52 and 0.02, respectively. This supports our decision to keep the sampling frequency for this farm type constant in our analyses at once per production cycle. This single sampling is assumed to take place at 17 weeks into the production cycle, which is approximately one week before distribution of the pullets to layer farms. A later sampling time is not possible since it takes around a week to obtain the test result.

In the absence of surveillance, most farms with an R_0_ value above 1 were located in the North of Israel. For SE, there was also clusters of farms with an R_0_ value above 1 further South (S3 and S4 Figs).

### The effect of the current surveillance programme on the between-farm R_0_

The current surveillance programme was predicted to reduce the between-farm R_0_ of SE below 1 for 98.2% of layer farms (S1 Fig). For ST, this is the case for all layer farms (S2 Fig). The highest between-farm R_0_ was 1.09 for SE and 0.45 for ST.

### Comparison of the general and risk-based surveillance programmes

To compare the different general and risk-based surveillance programmes, we looked at the yearly number of sampling events that resulted in R_0_<1 for all layer and pullet farms. For the general surveillance programme targeting both SE as well as ST (serovar scenario C), the sampling interval of layer farms had to be reduced from 15 weeks in the current programme to 13 weeks (Table 2) to reduce the R_0_ of both target serovars below 1 for all layer farms. This corresponds to an increase by approximately 15% of the yearly number of sampling events (Table 2). However, the risk-based surveillance programme could achieve the same level of disease control (R_0_<1) for both target serovars using a sampling rate that was 41% lower in comparison to the optimized general surveillance programme and 32% lower in comparison to the current surveillance programme (Table 3). This reduction was achieved by assigning layer farms in an area with a farm density below 2.4 farms per km^2^ (calculated using a circular area around the farm with a 3.91 km radius) to a low-risk group and the remainder of farms to a high-risk group (Fig 3) and using group specific sampling rates. The vast majority of layer farms was in the low-risk group (69%) and sampled every 34 weeks, which is much longer than the sampling interval of the general surveillance scheme. The high-risk group was sampled using the same interval as in the general surveillance scheme (13 wks). Most of the layer farms in the high-risk group were located in the North of Israel (Fig 4).

**Fig 3.**
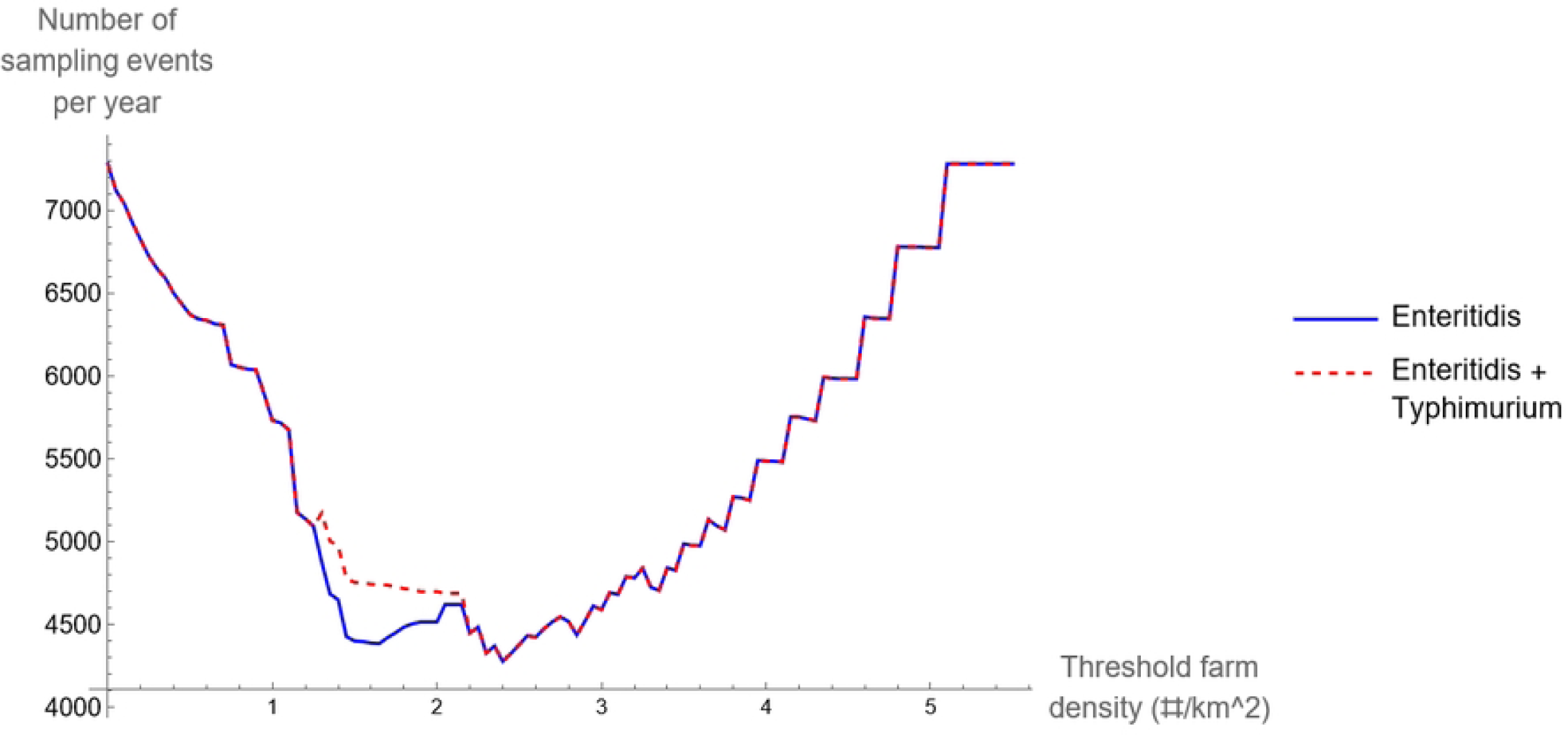
The yearly number of sampling events for the risk-based surveillance programme as a function of the local farm density (in a circular area of 4 km around a given farm), that was used to divide layer farms in Israel into groups with a low and a high risk of between-farm transmission of *Salmonella* Enteritidis alone or both Enteritidis and Typhimurium.

**Fig 4.**
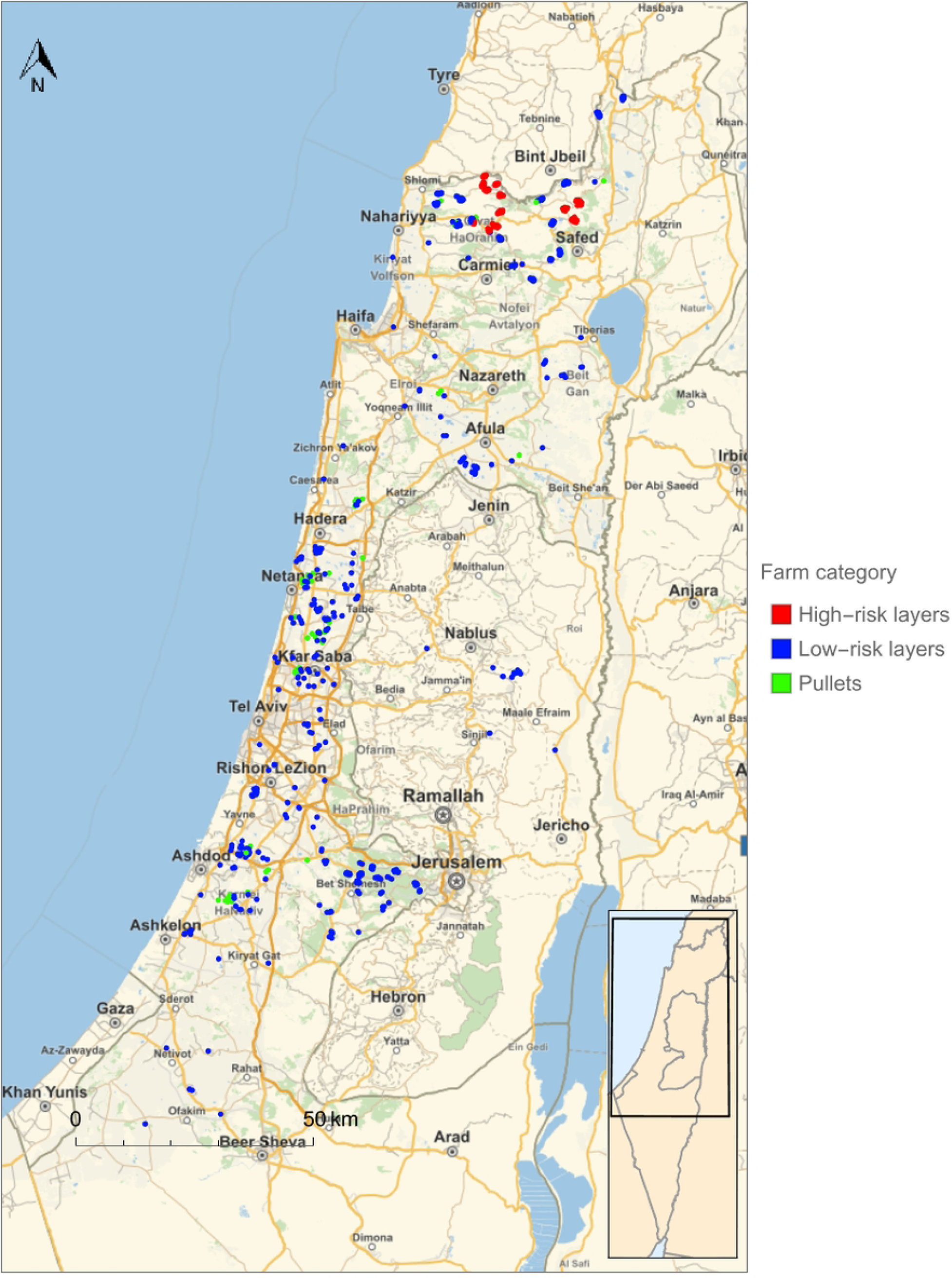
The division of layer farms in Israel into groups at low and high-risk of infection with *Salmonella* serovars Enteritidis and/or Typhimurium as a result of between-farm transmission.

**Table 2.** Characteristics of the current and optimized general surveillance programmes for *Salmonella* serovars Enteritidis and/or Typhimurium for pullet and layer farms in Israel.

| *Salmonella* serovar targeted | Sampling interval (wks) |  | Number of farms |  | Yearly number of sampling events | Highest between-farm R<sub>0</sub> |  |
| --- | --- | --- | --- | --- | --- | --- | --- |
|  | Pullet<sup>a</sup> | Layer | Pullet | Layer |  | Serovar Enteritidis | Serovar Typhimurium |
| *Current general surveillance programme* |  |  |  |  |  |  |  |
| Enteritidis and Typhimurium | 17 | 15 | 53 | 1782 | 6331 | 1.08 | 0.45 |
| *Optimized general surveillance programme* |  |  |  |  |  |  |  |
| Enteritidis | 17 | 13 | 53 | 1782 | 7281 | 0.998 | - |
| Typhimurium | 17 | 38 | 53 | 1782 | 2592 | - | 0.997 |
| Enteritidis and Typhimurium | 17 | 13 | 53 | 1782 | 7281 | 0.998 | 0.9 |

<sup>a)</sup> Pullet farms were always sampled one week prior to the distribution of pullets to layer farms.

**Table 3.** Characteristics of the optimized risk-based surveillance programmes for *Salmonella* serovars Enteritidis and/or Typhimurium for pullet and layer farms in Israel.

| *Salmonella*<br>serovar<br>targeted | Sampling<br>interval<br>pullets<sup>a</sup> | Sampling interval<br>layers |  | Number<br>of pullet<br>farms | Number of layer<br>farms |  | Yearly<br>number of<br>sampling<br>events | Highest between-farm R<sub>0</sub> |  |
| --- | --- | --- | --- | --- | --- | --- | --- | --- | --- |
|  |  | Low-risk<br>group | High-risk<br>group |  | Low-risk<br>group | High-risk<br>group |  | Serovar<br>Enteritidis | Serovar<br>Typhimurium |
| Enteritidis | 17 | 34 | 13 | 53 | 1215 | 567 | 4279 | 0.998 | - |
| Typhimurium | 17 | 52 | 38 | 53 | 1729 | 53 | 1955 | - | 0.997 |
| Enteritidis<br>and<br>Typhimurium | 17 | 34 | 13 | 53 | 1215 | 567 | 4279 | 0.998 | 0.90 |

<sup>a</sup>) Pullet farms were always sampled one week prior to the distribution of pullets to layer farms.

The comparison of the general and risk-based surveillance programmes targeting only one serovar (scenarios A and B) also showed that the risk-based programme is more efficient than the general programme. The surveillance programmes targeting SE also reduced the R_0_ of ST below 1, reflecting that the R_0_ of SE was higher than for ST for most layer farms. The optimum surveillance programmes targeting ST therefore provided sufficient disease control at much lower sampling rates than programmes targeting SE (Table 3). In addition, the optimum threshold density for dividing layer farms into low-risk and high-risk groups for SE transmission was relatively high (Fig 5), resulting in 97% of layer farms falling into the low-risk group with the lowest sampling rate. Most of the layer farms in the high-risk group were again located in the North of Israel (Fig 6).

**Fig 5.**
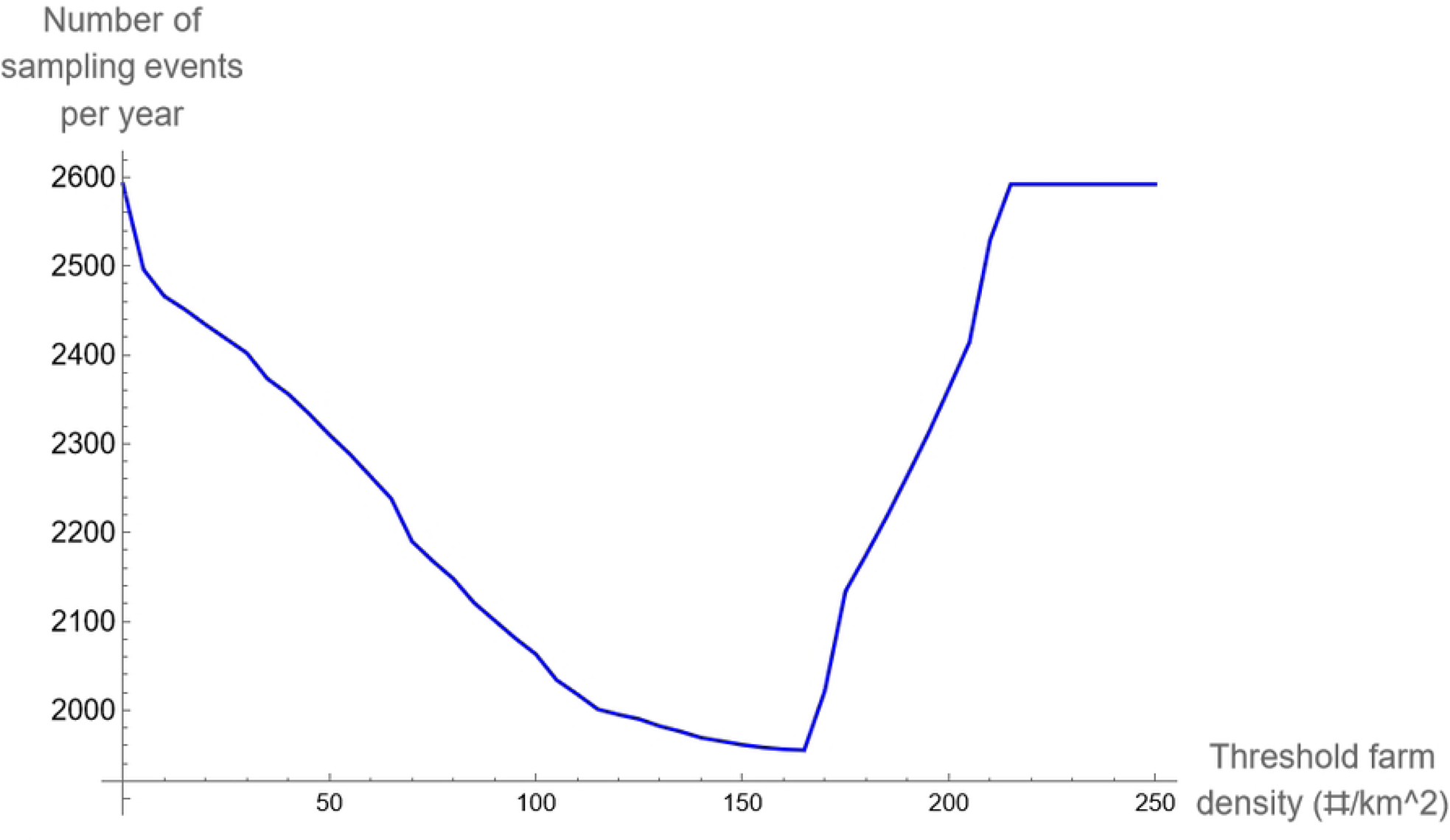
The yearly number of sampling events for the risk-based surveillance programme as a function of the local farm density (in a circular area of 300 m around a given farm), that was used to divide layer farms in Israel into groups with a low and a high risk of between-farm transmission of *Salmonella* Typhimurium.

**Fig 6.**
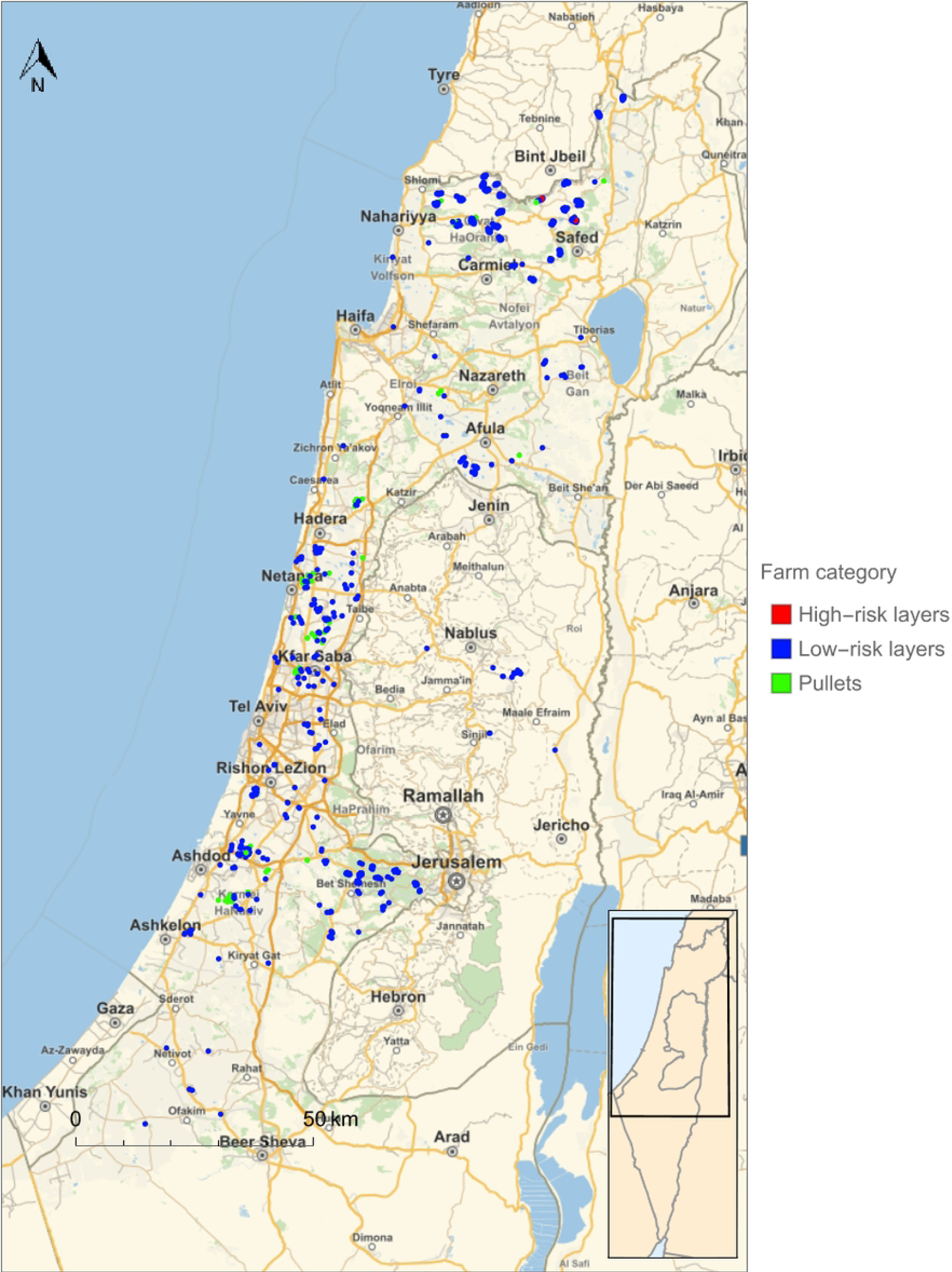
The division of layer farms in Israel into groups at low and high-risk of infection with *Salmonella* serovar Typhimurium as a result of between-farm transmission.

## Discussion

In this study, we determined the significance of local farm density as a risk factor for the infection of Israeli pullet and layer farms by SE and ST and we explored the shape of the transmission kernel that describes the relationship between the infection probability and the distance to an infected farm. The results show that the infection probability is significantly higher for both target serovars when the distance to an infected farm decreases below a certain serovar-specific threshold. Interestingly, we found that SE is transmitted over much longer distances than ST. These findings were subsequently used to design and evaluate a risk-based surveillance programme that determines the sampling frequency of layer farms based on the local density of farms. The results showed that a risk-based *Salmonella* surveillance programme for pullet and layer farms can provide the same level of disease control as a general surveillance programme, but using a much lower sampling effort. Surveillance programmes controlling SE also provided sufficient control of ST.

To our knowledge, three previous studies included the distance to the nearest farm (presumably of the same type) as a categorical variable in a risk factor analysis of *Salmonella* infections on layer farms. When defining a farm positive when infected by any *Salmonella* serovar, two of these studies[11, 12] showed that the infection probability was significantly higher for distances to nearest farm closer than 0.5 or 1 km. By contrast, the third study[10] showed a significant increase of the infection probability for distances to nearest farm larger than 1 km. The first two studies were however conducted in England and Italy, while the third was conducted in Nigeria, which may have a poultry industry that is less structured and less regulated with respect to animal health in comparison to the Israeli poultry sector. Although the results of the studies in England and Italy are not directly comparable to ours, too, since they included infections by any serovar, they seem more relevant for Israeli layer farms and are in broad agreement with our results for SE and ST. The English study also determined the effect of the distance to the nearest farm on the infection probability of layer farms by serovar SE in specific[11]. The results showed that the distance to the nearest farm was not a significant risk factor for infection by SE in England. We could not find any studies on ST in specific. It can be concluded that the few existing studies provide a mixed picture with some studies suggesting that a smaller between-farm distance may increase the infection probability, some studies suggesting the opposite and another study suggesting that there is no significant effect at all.

We found only one study that explored the relationship between the occurrence of layer farms in Iran that tested positive for any *Salmonella* serovar and the farm density in a 1 km^2^ area around a farm[13]. The results showed that all positive farms were located in areas with a relatively low farm density and therefore do not imply that a higher local farm density increases the infection probability by *Salmonella*.

A direct comparison of the results of our analyses with the results of the literature studies above is not possible for the following reasons. Firstly, instead of just looking at the distance to the nearest farm[10–12], our kernel functions for SE and ST were derived using a method that calculates the infection/escape probability of a farm at a given point in time using information on the distance to all other farms and their infection status at that time. Secondly, instead of treating distance as a categorical variable, our analysis allows the data to inform the shape of the transmission kernel. This in turn allowed us to derive the most relevant area for calculating the local farm density as risk factor from the fitted kernel function. Using a predetermined area for calculating the local farm density[13] decreases its relevance as risk factor, because the actual area within disease transmission is possible may be smaller or larger than the predetermined area. For these reasons, we think that the kernel method that we used in this study is better suited to analyse the effect of the distance to infected farms on the infection probability and explore the importance of local farm density as a risk factor for *Salmonella* transmission.

The results suggest that the radius of the area within which between-farm transmission is possible is much larger for SE (4 km) than for ST (0.3 km). This indicates that the processes underlying between-farm distance also differ between SE and ST. Two recent studies have reviewed the existing literature on risk factors for *Salmonella* infection of layer farms[17, 18]. Significant risk factors that may be associated with an increased probability of between-farm transmission of SE in specific were allowing visitors into the farm house[19], the presence of rodents, flies or wild birds in and around farm houses[11, 19] and the presence of dogs and cats on a farm itself or on contiguous farms[11]. One study also suggested that an open ventilation system increased the risk of infection by SE compared to a closed system[20], which implies that *Salmonella* may also be dispersed via airborne particles, although it is unknown at what spatial scale this may take place. We did not find studies on risk factors that may be associated with between-farm transmission of ST in specific. On the contrary, many studies (only) analysed the significance of risk factors for the infection of layer farms by any *Salmonella* serovar. Some of these studies reported that the presence of trucks near the entrance of farm houses and air inlets (e.g. for feed delivery or the collection of eggs and dead birds) increased the risk of *Salmonella* infection[11, 21, 22]. Our study suggests that the processes underlying between-farm transmission of ST operate over short distances of a few 100 meters and may therefore be explained by the movement of biological and mechanical vectors such as rodents, flies and wild birds between farms. The transmission of SE occurs over much longer distances of several kilometers and may be explained by the movement of vehicles between-farms. It is however not clear why SE would be able to spread via vehicles, but ST not. Perhaps SE is able to survive in the environment for a longer period of time then ST.

Even in the virtual absence of between-farm transmission due to an intensive surveillance programme, pullet and layer farms can therefore still become infected by *Salmonella* via introduction from a local environmental source. *Salmonella* serovars can use a broad range of host species including mice and rats that may be present in the environment of farms[23]. In addition, *Salmonella* may survive for long periods of time outside host species, *e.g.* in soil[23] and to a lesser extent also in water[24]. Based on the results of our study, we estimated that the probability of a flock of laying hens becoming infected from a local environmental source amounts to 1.2% for SE and 1.8% for ST (derivation not shown). In order to further reduce these percentages, current biosecurity protocols and/or the compliance with these protocols have to be improved. Biosecurity measures that reduce the likelihood of introduction from a local environmental source may also reduce the probability of between-farm transmission.

In this study, we optimized surveillance programmes using the requirement that the between-farm R_0_ for target serovars SE and ST was below 1 for all pullet and layer farms. In addition, we assumed a maximum sampling interval for layer farms of 52 weeks. Clearly, somewhat different results could be expected when other or additional criteria would be used such as a lower/higher maximum value for the between-farm R_0_ or a shorter maximum sampling interval (ultimately reducing the number of human Salmonellosis cases).

One of the limitations of this study is that the we only included pullet and layer farms in our analysis, while *Salmonella* may also be present on other types of poultry farms (for example broiler, turkey and duck farms), pig farms and cattle farms. A recent European Union report showed that the prevalence of SE was highest on layer farms, while the prevalence of ST was highest on pig and cattle farms[2]. It is not clear how transmission of SE and ST between the farm types included in our study and other farm types would change the importance of the local density of pullet and layer farms as a risk factor for infection by these serovars. However, the number of contacts between different farm types may be limited since they may not be part of the same supply chain and similar farm types may be concentrated in specific regions.

Another limitation is the fact that, in our analyses, we did not account for the sensitivity of the dust and drag swab sampling methods and the sensitivity of the diagnostic laboratory tests for identifying SE and ST. The actual prevalence and persistence of these serovars on pullet and layer farms may be higher than the longitudinal dataset shows due to false negatives. It is however unclear if and how this would have affected the shape of the transmission kernels for SE and ST. We also did not account for the sampling and test sensitivity when evaluating the different surveillance programmes. If an SE or ST infection is not detected during a sampling event, this means that the farm will not be culled and remains infectious for a longer period of time, which in turn increases the between-farm R_0_ for this farm. The reduction in between-farm transmission achieved by a surveillance programme may therefore be less than our analyses predicted. However, our analysis also showed that most farms have a between-farm R_0_ that is much lower than one for the current and optimized surveillance programmes (S1 and S2 Figs). Therefore, the between-farm R_0_ of many farms may still be lower than 1 even if corrected for the possible underdetection.

Finally, in this study, we evaluated and designed surveillance strategies based on the between-farm R_0_. Information on the between-farm R_0_ is however not sufficient to predict the temporal change in the disease prevalence and the number of infected farms in time due to a given surveillance programme. This would require an actual stochastic simulation model that uses the transmission kernel to predict the probability of between-farm transmission. In addition, criteria for the optimization of surveillance strategies that weigh the costs of a surveillance programme against the benefits in terms of e.g. human quality of life years gained (for example the incremental cost-effectiveness ratio) would be needed.

We conclude that the local density of pullet and layer farms can be a significant risk factor for the infection of these farms by SE and ST. The relationship between the infection probability of pullet and layer farms by these serovars and the distance to an infected farm can be described using a transmission kernel that assumes a threshold distance above which between-farm transmission is unlikely. *Salmonella* surveillance programmes for pullet and layer farms that account for the local farm density as a risk factor for infection by SE and ST may provide the same level of disease control as general surveillance programmes, while substantially reducing the required sampling effort.

## Acknowledgements

We thank Gert-Jan Boender from Wageningen Bioveterinary Research in the Netherlands for his insights and advice about the shape of the transmission kernel in our analysis.

## Supporting information

**S1 Fig.** The distribution of the predicted between-farm R_0_ of *Salmonella* Enteritidis for pullet and layer farms in Israel in the absence of surveillance and under the current surveillance programme (see main text).

**S2 Fig.** The distribution of the predicted between-farm R_0_ of *Salmonella* Typhimurium for pullet and layer farms in Israel in the absence of surveillance and under the current surveillance programme (see main text).

**S3 Fig.** The predicted between-farm R_0_ for *Salmonella* Enteritidis in the absence of a surveillance programme for pullet and layer farms in Israel.

**S4 Fig.** The predicted between-farm R_0_ for *Salmonella* Typhimurium in the absence of a surveillance programme for pullet and layer farms in Israel.

